# Elemental content of a host-parasite relationship in the threespine stickleback

**DOI:** 10.1101/2022.05.04.490668

**Authors:** Megan Braat, Rita L. Grunberg, Daniel I. Bolnick

## Abstract

Parasite infections are ubiquitous and their effects on hosts may play a role in ecosystem processes. Ecological stoichiometry provides a framework to study linkages between consumers and ecosystem process, but the stoichiometric traits of host-parasite associations are rarely quantified. Specifically, whether parasites’ elemental ratios closely resemble those of their host or if infection is related to host stoichiometry remains less known. To answer such questions, we measured the elemental content (%C, %N, and %P) and ratios (C:N, C:P, and N:P) of parasitized and unparasitized *Gasterosteus aculeatus* (three-spined stickleback) and their cestode parasite, *Schistocephalus solidus*. Host and parasite elemental content were distinct from each other, and parasites were generally higher in %C and lower in %N and %P. Parasite infections were related to some elemental ratios, specifically C:N, with more intense parasite infections corresponding to hosts with lower C:N ratio. Parasite stoichiometry was independent of their host and there was no relationship between host and parasite stoichiometry. Instead, parasite body mass and parasite density were important drivers of parasite stoichiometry where larger parasites had lower %C, %N, and %P,. Overall, these potential effects of parasite infections on host stoichiometry along with parasites’ distinct elemental compositions suggest parasites may further contribute to how hosts store and cycle nutrients.

## Introduction

Consumers are important components to nutrient cycling in aquatic ecosystems (Atkinson et al., 2017). For example, fish play a role in nutrient dynamics in aquatic systems, both through nutrient storage and excretion (Vanni, 2002; Vanni et al., 2013; Vanni and McIntrye 2016). Variation in nutrient cycling between species can broadly be explained by differences in the elemental composition of species (Elser et al., 1998 Ecosystems, Elser and Urabe, Ecology). Ecological stoichiometry (ES) provides a mechanistic framework to link differences in the relative quantities and balance of elements between consumers and resources to ecosystem processes (Sterner & Elser, 2002). Within this framework, heterotrophic consumers, such as fish, are thought to maintain a relatively constant elemental composition (Vanni et al., 2002), and thus focus has been on describing among-species variation in organismal stoichiometry. However, it is now clear that within-species variation in elemental composition is substantial and sometimes exceeds variation among species (El-Sabaawi et al., 2012, Lemmen et al., 2019). This stoichiometric variation, in turn, can modify nutrient flows through ecosystems to alter productivity and community structure. Consequently, a better understanding of the drivers of intraspecific variation in organismal stoichiometry is necessary to further link individuals to ecosystem processes.

Within fish species, organismal stoichiometry is determined by a myriad of factors that interact with physiology, such as environmental variation, sex differences, ontogeny, and genetics (Boros et al., 2015; El-Sawbaawi et al., 2016, 2012, Leal et al., 2017; Mozsar et al.., 2019). Disease is another potential driver of fish stoichiometry but has received relatively little attention. Fish are commonly infected with a diversity of parasite species that impact fish physiology and behavior (Barber et al., 2017), but the potential effects of infection on fish stoichiometry and excretion are not widely known. Evidence from snail-trematode systems suggest parasite infections contribute to variation in host stoichiometry and excretion chemistry (Bernot 2013, Chodkowski and Bernot 2017; Mischeler et al., 2016). Parasite infections can increase nitrogen excretion by their snail hosts when parasites require more carbon relative to nitrogen from their host (Mischeler et al., 2016). Nevertheless, the consequences of infection will ultimately be contingent on the host-parasite association, and the corresponding organismal stoichiometry of the parasite and host. Currently, there is a paucity of data on the organismal stoichiometry of parasitic species (but see Paseka and Grunberg 2019), and stoichiometric data on paired host-parasite associations from fish are rare. This is a critical knowledge gap, given that consumers, here fish and parasites, vary widely both within and among species, and variation in organismal stoichiometry can affect ecosystem processes (El-Sawbaawi et al 2016).

To understand linkages between parasite and host stoichiometry, we asked three questions. First, do parasites’ elemental content match that of their hosts, or do they have different elemental traits? Second, are host and parasite stoichiometry correlated, such that hosts with higher C:N ratios (for instance) have parasites with correspondingly higher C:N? Lastly, are parasite infections related to differences in host stoichiometry? Such relationships might arise because parasites induce immune responses that cause stoichiometric changes, or because the parasite disproportionately extracts certain nutrients from its host. We addressed these questions by collecting samples of three-spine stickleback (*Gasterosteus aculeatus*) from a wild population, contrasting fish with or without natural infections by the cestode parasite *Schistocephalus solidus*. We measured the C, N, and P content and their respective ratios for both infected and uninfected stickleback, and for the cestode when present. The three-spined stickleback (*Gasterosteus aculeatus*) exists in both freshwater and marine ecosystems and is host to a diversity of parasites (Bolnick et al., 2020). Our study focused on the parasite S. solidus which infect stickleback when the fish consume infected copepods. The parasite exits the zooplankton and burrows through the gut to establish a long-lasting infection in the body cavity, where the parasite gains nutrients from the host’s circulatory system (though the details are poorly known). This parasite offers several advantages for this study. Foremost, this parasite has a large body mass, which allows us to quantify C, N, P for an individual parasite, and pair these data with an individual host. Second, by living in the body cavity rather than the gut, the parasite must gain its resources from the host, rather than directly from the host’s food in the intestines. Further, *Schistocephalus solidus* infections are known to affect host physiology (Barber & Scharsack, 2010), immunity (Scharsack et al., 2004; Lohman et al., 2017; Weber et al., 2017), morphology (Dingemanse et al., 2009)(Ness & Foster, 1999), and behavior (Talarico et al., 2017), which are likely to impact how hosts allocate resources.

## Methods

### Stickleback collection and necropsy

We collected three-spined stickleback from Merrill Lake on northern Vancouver Island in British Columbia, Canada on October 5, 2020. A total of 95 fish were collected from the lake using a dipnet and were stored in a freezer until dissection. From this sample, we chose 30 infected individuals and 30 uninfected individuals for chemical and parasitological analysis. Stickleback were sampled with approval from the University of Connecticut Institutional Animal Care and Use Committee (IACUC; protocol A18-008), and with a scientific fish collection permit from the British Columbia Ministry of the Environment (permit NA20-600806).

We thawed individual fish and measured standard length (snout to caudal peduncle) and wet mass for each specimen, and subsequently dissected and sexed each fish through visual inspection of gonads. We quantified the presence, intensity (number of cestodes), and mass of *S. solidus*. Tapeworms were retained for chemical analyses.

### CNP analysis

After dissections, we dried each fish and its associated parasites (if infected) by storing samples in a drying oven at 60 °C for 7 days, and then weighing each sample on a microbalance to measure total dry mass. Later, we homogenized each sample into a fine powder with a mortar and pestle and stored samples in a desiccator until CNP analysis. We used 1-2 mg of each homogenized sample of *G. aculeatus* and *S. solidus* for element analysis. Samples were sent to the University of Georgia Center for Applied Isotope Studies Stable Isotope Ecology Laboratory to quantify total percent C, N, and P, and their respective ratios.

### Statistical Analysis

The overarching goal of our study was to describe differences in host and parasite organismal stoichiometry and evaluate whether differences in host stoichiometry were related to parasite infections. Our first question was whether hosts and parasites differ in their elemental composition. *S*.*solidus* constitutes a large proportion of host biomass, so if they disproportionately uptake certain elements from the host, this could change host stoichiometry as well. To determine whether stickleback and *S. solidus* have distinct elemental compositions, we used paired t-tests to compare the species’ means for %C, %N, and %P and their respective ratios, C:N, C:P, and N:P. In addition, we also

Prior studies have found significant variation in stoichiometry among individual fish (Vanni et al., 2002; Sterner and George, 2000; Mozsár et al. 2019), which may lead to stoichiometric variation among parasites. For example, an increase in %N across hosts could be related to an increase in %N of its parasite. To test this hypothesis, we calculated Pearson correlations (and associated P-values) between paired host and parasite elemental contents, and stoichiometric ratios.

Host stoichiometry could be related to parasite infections if parasites alter host stoichiometry or if hosts with certain stoichiometry ratios are more likely to be infected. To test this, we estimated linear models in which host elemental content or ratios (C:N, C:P or N:P) depends on host infection status (infected or uninfected) after correcting for effects of host body mass. In a follow up analysis, we tested the impact of parasite infection intensity (# parasites per host) on infected host stoichiometry by estimating another linear model in which host elemental content or ratios depends on parasite infection intensity and host body mass.

Using a multiple regression, we also asked if parasite elemental composition was related to both parasite body mass and parasite density (# parasites/ dry mass of host). Within and across parasite species, body mass is negatively related to phosphorus content (Paseka and Grunberg, 2019), because of its relationships with growth rates (i.e., growth rate hypothesis, Sterner and Elser 2003. Relationships between parasite elemental content and parasite density may also arise if parasite density affects parasite growth rates. Statistical analyses and graphics were produced in R studio ver. 1.4.1106 (R Core Team, 2020).

## Results

In total, we analyzed the elemental content of 25 uninfected fish, 27 infected fish, and 26 parasites (one per infected fish; one parasite was too small for elemental analysis). The parasite, *S. solidus*, on average constituted 22.4 +/- 3.1% SE of total host biomass (i.e., dry mass of S. solidus + G. aculeatus). The parasite’s maximum contribution to total host biomass was 58.6%, and its minimum was less than 1%.

### Elemental composition of the parasite of *S. solidus* and its stickleback host

The parasite *Schistocephalus solidus* had a distinct elemental composition from their stickleback host (Figure 1). In general, *S. solidus* had a higher %C content than their host (mean % C: 45.9% and 39.2%, respectively, paired t-value = 5.621, p < 0.0001). However, S. solidus were lower in %N (mean % N: 7.7% and 10.5%, respectively, paired t-value = -7.819, p < 0.0001) and %P relative to their host (mean % P: 1.6% and 4.4%, respectively, paired t-value = -12.481, p < 0.0001). *S. solidus* also had higher C:N, C:P, and N:P element ratios compared to their hosts (t-tests, p < 0.0001).

**Figure 1.**
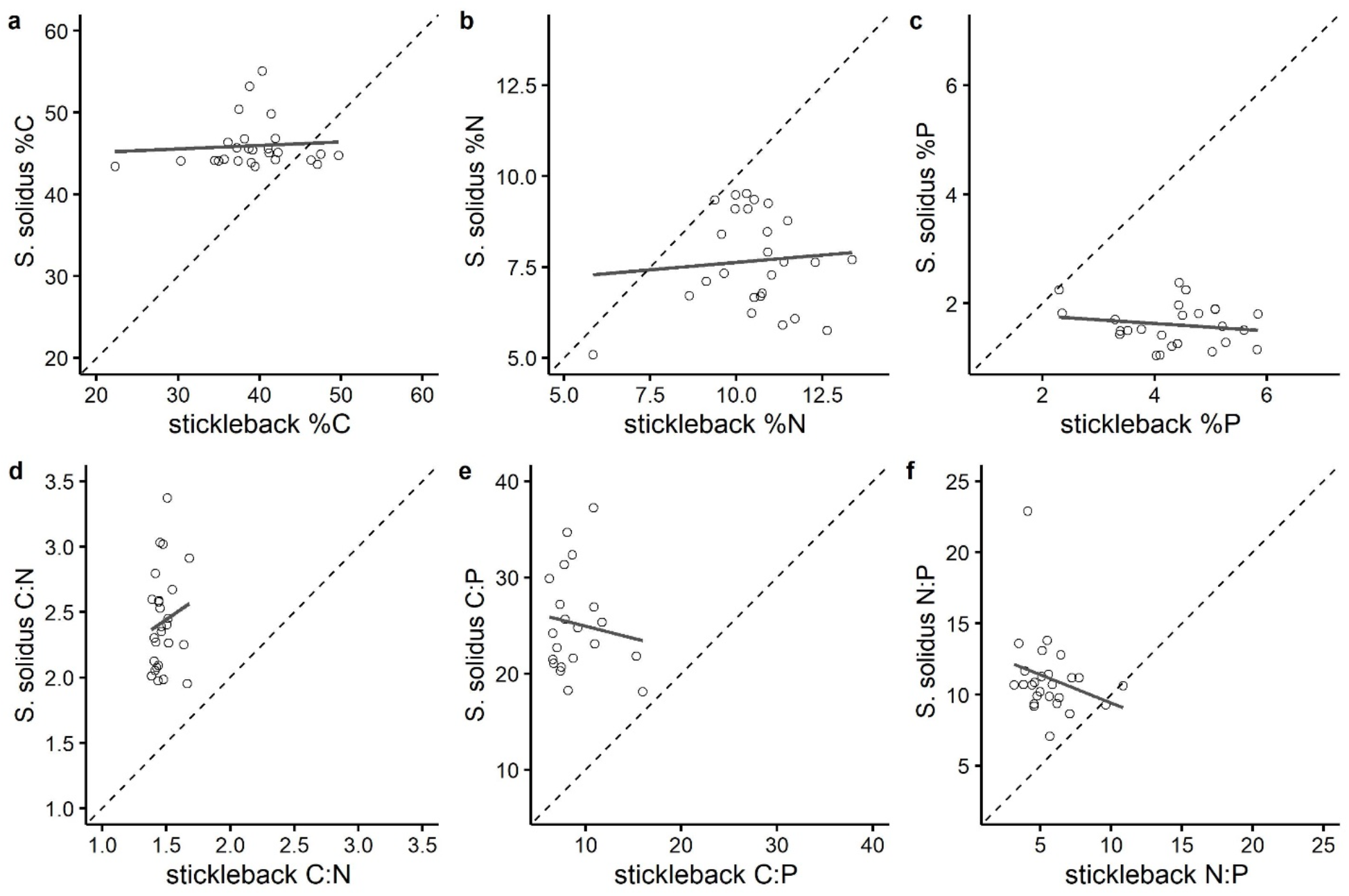
*Schistocephalus* and stickleback elemental composition were not correlated across host-parasite pairs (p >0.05). Instead, parasites appear to retain a relatively constant elemental composition that is independent of, and different from their host resource. A dashed line is drawn from the origin to show a directly proportional relationship, and the grey solid line is a trend line.

Host and parasite elemental content were not correlated across samples (all correlations p > 0.05, Figure 1). Generally, correlations between parasite and host %C (r = 0.07, p =0.698), %N (r = 0.08, p =0.669), and %P (r = -0.26, p=0.196) were weak and not significant. Similarly, there was no evidence of a relationship between parasite and host stoichiometric ratios for C:N (r = 0.14, p=0.483), C:P (r = -0.307, p =0.126), and N:P (r = -0.249, p=0.219). Thus, elemental variation among individual hosts did not impact the elemental composition of their respective parasites.

### Relationship between parasite infection and host stoichiometry

Differences in elemental composition between host and parasite might reflect a zero-sum process in which certain elements are disproportionately retained by the parasite and lost by the host. The result would be no net change in the elemental composition of the host-parasite combination. Alternatively, infection may alter the host’s intake or excretion of elements, changing the host-parasite elemental composition. Our data support the latter scenario: uninfected fish differed from the host-parasite combination. The host – parasite phenotype was significantly higher in %C, and lower in %N and %P relative to the infected host tissue alone (p > 0.001, Figure 2).

**Figure 2.**
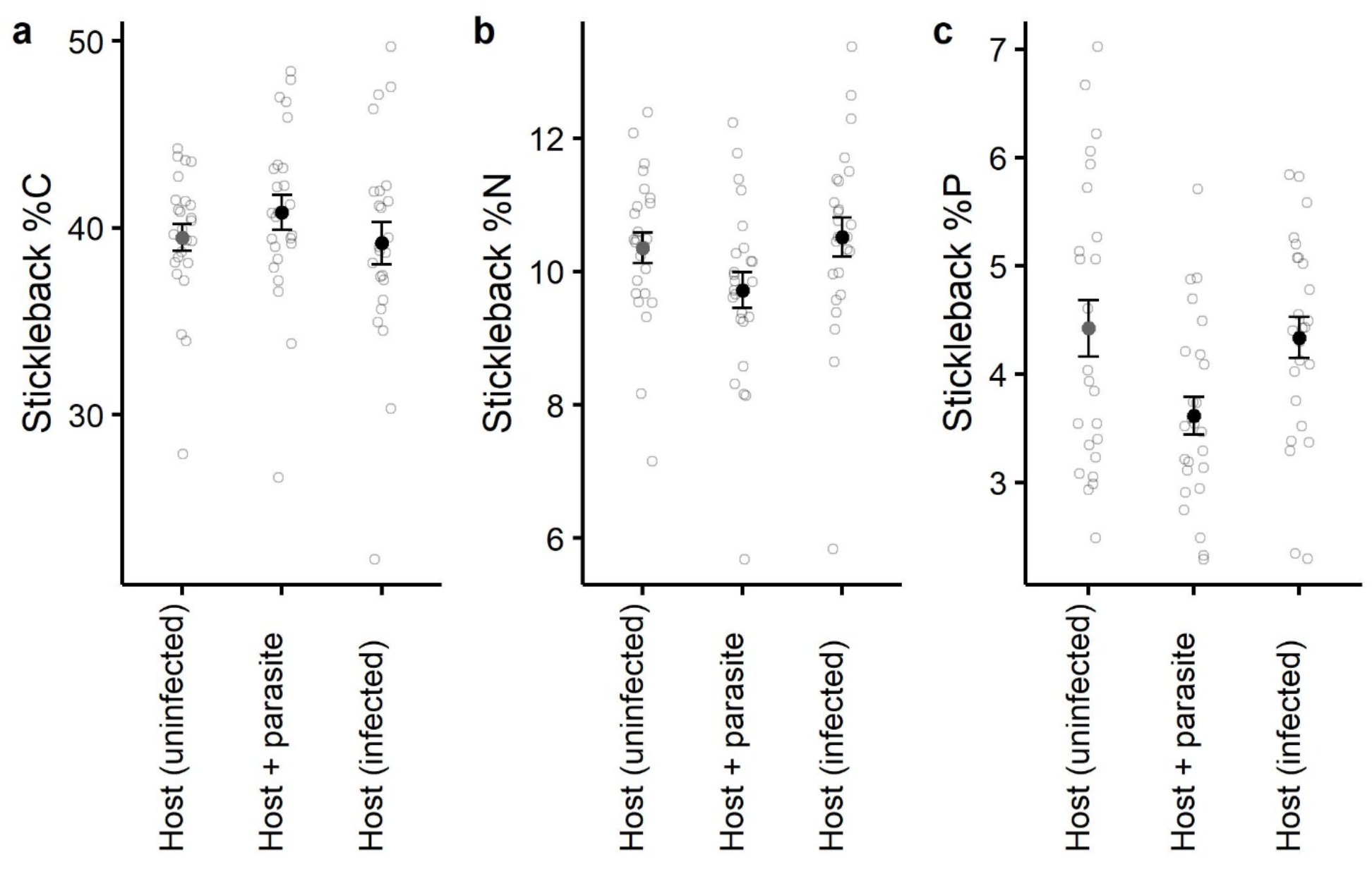
The host infection phenotype differed when considering the host + parasite as a collective unit relative to the infected host tissue alone and uninfected host tissue. The host + parasite phenotype was significantly higher in %C (a), and lower in %N (b) and %P (c) when compared to the infected host tissue alone.

This difference between infected versus uninfected fish persists when we consider host tissue alone (e.g., excluding parasite elemental composition). Using a linear model that accounts for host body mass as a covariate, infected hosts were lower in C:N than uninfected hosts (p = 0.03, R^2^= 0.09, Figure S1). This negative relationship between C:N and infection was also apparent when including parasite infection intensity in our model, which revealed that hosts carrying a larger number of tapeworms had lower C:N (p = 0.04, R^2^= 0.10, Figure 3). However, host total %C and %N were not related to parasite infection status or intensity (p > 0.05). All other relationships between host infection and host elemental ratios were not significant (p > 0.05, Figure 2).

**Figure 3.**
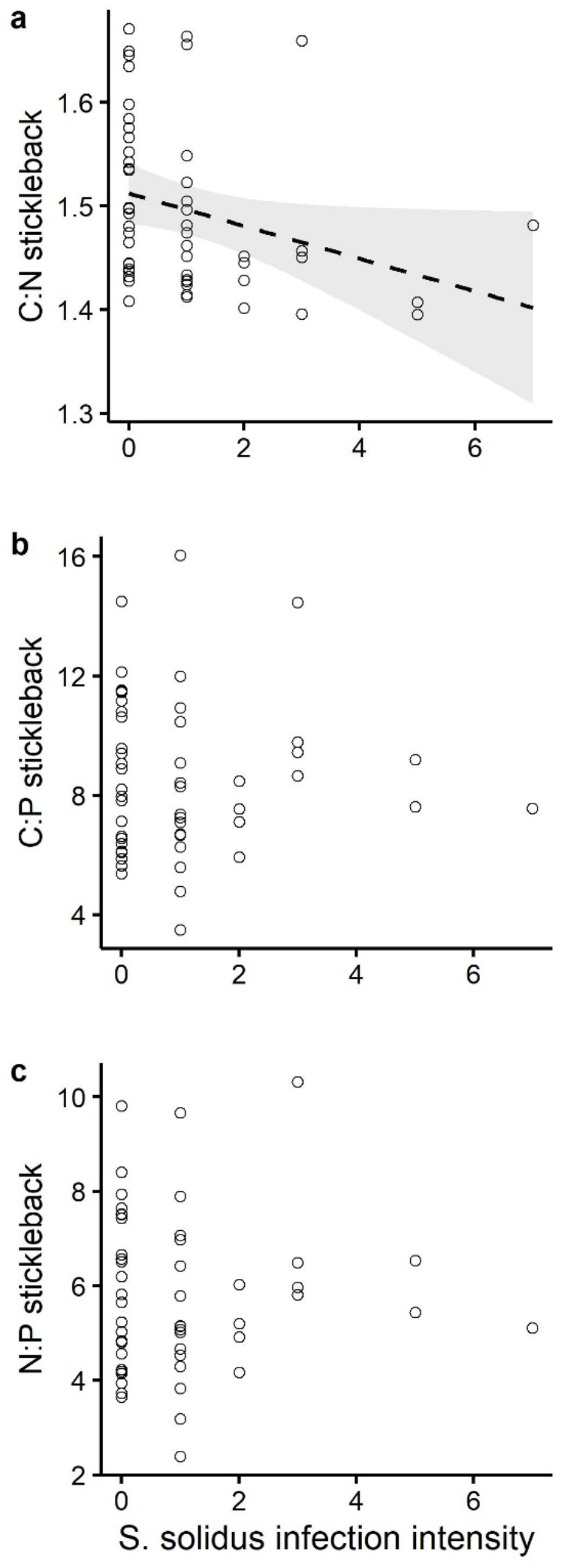
Host C:N (a) was related to parasite infection intensity after accounting for host body mass as a covariate. However, host C:P (b) and N:P (b) were not related to parasite infection intensity. Plotted are the partial residuals of the linear model that account for the effects of host standard length on host stoichiometry. Dashed line are the estimated model fits and grey shaded area is the 95% confidence interval for the one significant trend.

### Parasite stoichiometry

Differences in parasite elemental content were generally driven by parasite body mass and parasite density (Figure 4). As *S. solidus* body mass increased total carbon content decreased (slope = – 2.23, p = 0.003; Figure 4a), but infection intensity (e.g., crowding) was not related to differences in parasite carbon content (slope = 0.63, p = 0.531; Figure 4b). Together, parasite body mass and intensity explained 53% of variation in parasite %N and 52% of variation in %P. *S. solidus* nitrogen content decreased with their body mass (slope = – 0.95, p = 0.016; Figure 4b) and increased with their density (slope = 1.43. p =0.014, Figure 3c). Similarly, phosphorus content also decreased with S. solidus body mass (slope = – 0.27, p = 0.009; Figure 4d) and increase with density (slope = 0.32, p =0.03, Figure 4e).

**Figure 4.**
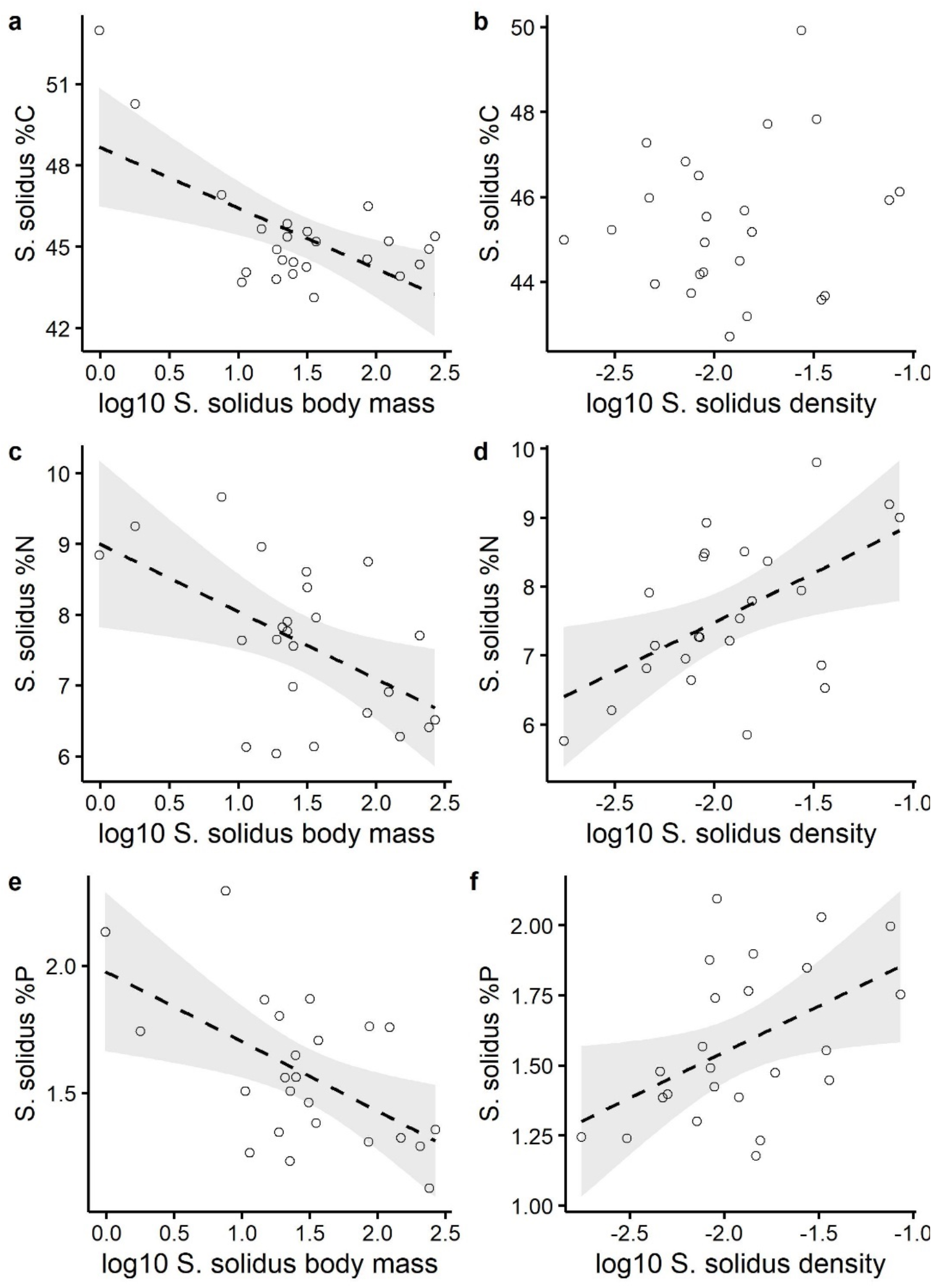
Parasite %C, %N, and %P decreased with parasite body mass and increased with parasite density (# parasite/host body mass (mg)). Plotted are the partial residuals of the statistical model for parasite %C (a-b), %N (c-d), and %P (e-f). Dashed lines are the estimated model fits and grey shaded area is the 95% confidence interval.

Relationships between parasite body mass, parasite density, and parasite elemental ratios were more variable (Figure S2). The C:N ratio of parasites generally decreased with parasite density (slope = -0.47, p=0.008) and showed weak evidence for a positive relationship with parasite body mass (slope =0.21, p=0.07). Parasite C:P increased with parasite mass (slope =-3.54, p=0.03) and decreased with parasite density (slope =-5.99, p=0.01). However, variation in parasite N:P was not related parasite mass (slope =0.51, p=0.34) and density (slope = -0.35, p=0.66).

## Discussion

Parasites have the potential to interact with ecosystem processes by shaping how hosts store and cycle nutrients (Mischeler et al 2016; Borer et al 2022). The effects of parasites on host stoichiometry should be driven in part by the relative differences in the elemental content between parasites and their hosts. However, few studies have tested for stoichiometric differences between, or correlations among, host and parasites, or how infection changes host stoichiometry (Chodkowski and Bernot 2017, Mischeler et al., 2016). Within the stickleback – *Schistocephalus* system, we show that parasite infections are related to differences in host C:N, but not C:P or N:P. Specifically, hosts with lower C:N ratios tended to have more parasites. In addition, parasite and host stoichiometry were not related across samples, so parasite stoichiometry appears to be independent of their host. Instead, differences in parasite stoichiometry was primarily driven by parasite body mass and parasite density. The observed differences between host and parasite stoichiometry, and between infected versus uninfected hosts, mean that parasitism contributes to variation in how hosts store and cycle nutrients in aquatic ecosystems.

### Parasite stoichiometry differs from host stoichiometry

Host and parasite elemental content and stoichiometric ratios were distinct from each other. As expected, sticklebacks were richer in phosphorus, which is related to their high investment in bony structures (El-Sabaawi, 2016), and nitrogen. On the other hand, *S. solidus* had a higher carbon content than their host. Parasites like *S. solidus*, which have long-lived larval stages and then fast maturation within their definitive hosts, may be expected to have greater levels of C-rich carbohydrates that are associated with maintenance and storage. This tendency to find C-rich parasites relative to their hosts has also been demonstrated in larval trematodes infecting snails (Bernot 2013, Mischler et al 2016). Differences in host and parasite stoichiometry reflects differences in their demands of certain elements and can contribute to variation in host cycling of nutrients (Mischler et al 2016). While we did not measure host excretion, we predict the higher carbon demands of S. solidus may shift host excretion towards a higher C:N concentration due to a greater assimilation of carbon by the parasite (Bernot 2013, Mischler et al 2016). Ultimately, variation in parasitism among hosts along with differences in host and parasite stoichiometry has the potential to shift nutrient availability and ecosystem processes (Bernot and Poulin, 2018).

The elemental composition of the parasite, *Schistocephalus solidus*, was independent of their host resource. Thus, hosts with higher concentration of an element do not necessarily have parasites with higher concentrations of that element. This could indicate that *S. solidus* selectively exploits its host for certain nutrients, maintaining an internal homeostasis. A general assumption is that some parasites consume bulk-goods from their hosts and thus parasites would have an enriched stable isotope value level relative to their host resource (Ben-David & Flaherty 2012; Behrmann-Godel & Yohannes 2015). However, Schistocephalus is frequently depleted in Nitrogen relative to their fish resource, so they are in effect feeding on a lower trophic level than their host (Pinnegar et al 2001, Power and Klein 2004, Eloranta et al 2015). Consequently, parasites that are capable of selective feeding within their host are less likely to experience changes in stoichiometry that are correlated with their host.

An important caveat is that we used whole-organism elemental analyses, but nutrients are distributed non-uniformly through the body (e.g., Phosphorous is enriched in bones). Thus, the tapeworms may deviate from elemental compositions of their host as a whole, but conform more closely to the composition of the particular host tissue(s) or blood supply from which they obtain nutrients (currently not clearly known).

Instead of covarying with host elemental composition, the cestodes’ C:N:P elemental content varied as a function of parasite body mass and density. The mechanisms underlying these relationships in parasites are not full known, but several hypotheses are plausible. First, parasites are likely exhibiting ontogenetic shifts in nutrient requirements as they transition from a fast-growing strategy, towards nutrient storage in preparation for breeding in their terminal hosts (birds, from whom they derive few if any nutrients). The negative relationship between parasite body mass and %P aligns with the growth rate hypothesis (Elser et al., 1996). Generally, rapidly growing organisms will be high in P since they will contain more P-rich ribosomal RNA, which is associated with protein synthesis. Consequently, P-content should be positively related to both growth rate and RNA content, and negatively related to body size (Peter, 1983) in organisms lacking P-rich bone structures. Second, as parasite density increases, they may compete as they grow, imposing more severe demands and alter their allocation to growth when resources are more limited. This potential competitive effect is consistent with the effect of both parasite mass and infection intensity.

### Infection is associated with host stoichiometry

Host C:N ratios were related to parasite infection status in our stickleback population, with infected individuals having lower C:N ratios than uninfected. This could be the result of a host’s coping strategy to infection. Hosts could be consuming more of their excess protein to compensate for the behavioral, morphological, and physiological changes in the host induced by the parasite directly, or by the host’s immune response to the parasite. Alternatively, *S. solidus* could be directly outcompeting *G. aculeatus* for its resources, specifically carbon. This nutrient deficit within the host could impact host immunity and the degree to which the parasite changes its hosts physiology. The latter scenario, however, would lead to a zero-sum redistribution of elements, which is inconsistent with our observation that infected fish (including their parasite) have different elemental composition than uninfected fish.

Stickleback carrying more cestodes (higher infection intensity) had lower C:N ratios. Since *S. solidus* have higher C:N ratios than their host, higher infection intensity apparently depletes their host of more carbon. This inference should be confirmed with future experimental infections, which we predict will show high host C:N ratios prior to infection, then gradually decreasing C:N ratios as *S*.*solidus* grows and depletes *G. aculeatus* of its carbon. Past research indicates that C:N ratios decrease as parasites reach maturity in other host species (Paseka & Grunberg, 2019). C:N ratios of *S. solidus* in our sample decreased as mass of *S. solidus* increased. This could indicate that larger cestodes were closer to reaching their reproductive stage and preparing to be transferred to their final avian host.

Because this is an observational study of wild-caught fish, we cannot currently exclude an important alternative explanation for our result: pre-existing stoichiometric variation might be associated with risk of cestode exposure. Stickleback are well known to exhibit among-individual variation in diet, within populations (Matthews et al., 2010; Araújo et al., 2008; Snowberg et al., 2015). Some individuals preferentially forage on benthic invertebrates, whereas others preferentially feed on limnetic zooplankton, including the copepods that are the first host for *S*.*solidus*. Diet differences thus contribute to predictable differences in infection risk among individuals within a given lake (Stutz et al.). Within-population diet variation generates variance in stable isotope ratios of Carbon, and Nitrogen, but we do not currently know whether diet variation also affects C:N:P stoichiometric ratios. If so, limnetic-feeding fish (with corresponding C:N:P ratios) might be more likely to be exposed to and infected by *S*.*solidus*, which could generate the correlations documented here.

Finally, all samples studied within our population had successfully survived up until the time of capture, which indicates a potential for survival bias. Individuals who are more tolerant of infection are more likely to be sampled in wild collections. Hypothetically, if stoichiometric ratios covary with tolerance, we might observe differences between surviving infected fish, compared to a typical uninfected fish.

### Conclusions

By comparing C,N,P composition of wild-caught individual fish and their individual parasites, we provide observational, correlational evidence that tapeworm infection is related to host stoichiometry. Conversely, the parasite stoichiometry is surprisingly independent of the host’s elemental composition. The parasite and host differ in average composition, and do not covary. These findings are significant because the parasite comprises a large fraction of the stickleback biomass in some populations, where infection prevalence can reach nearly 100% and over a quarter of fish biomass. Such large changes in stoichiometry of an abundant fish (often one of only a few fish species in north temperate coastal lakes), is expected to appreciably alter sticklebacks’ contributions to nutrient cycling. Prior studies from other fish have found that fish nutrient excretion can have substantial effects on ecosystem and community dynamics.

Therefore, parasitism has the potential to impart large changes in nutrient cycling in ecosystems, and warrant further investigation.

## Data accessibility

The data and code that support the findings of this study are available through Zenodo at https://zenodo.org/badge/latestdoi/484448578

## Acknowledgments

This work was funded by the University of Connecticut (start-up for DIB) and by the NIH 1R01AI123659-01A1 (to DIB).

---

**Figure S1.**
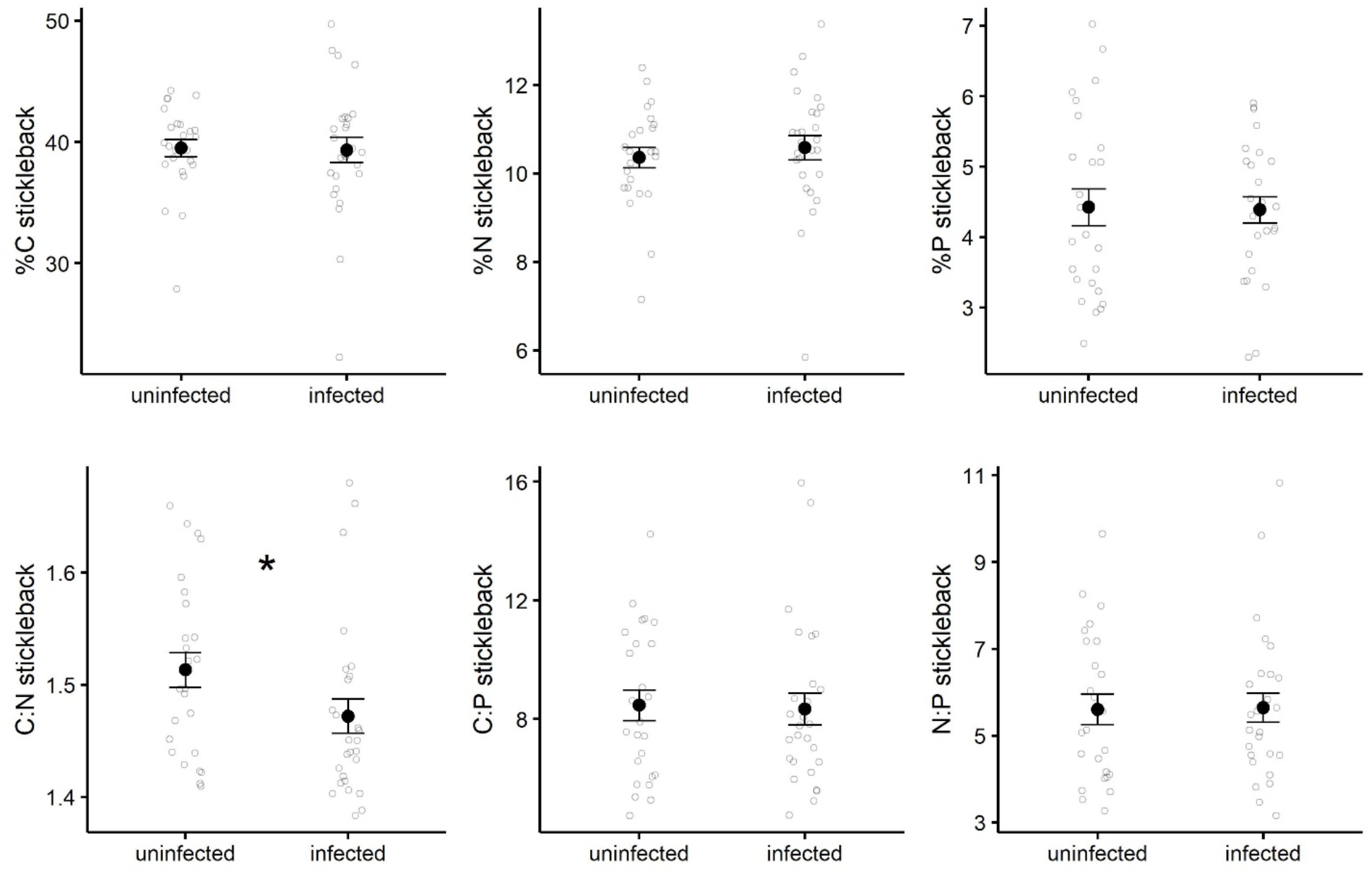
Differences in C, N and P, and their respective ratios in uninfected and infected stickleback. Host C:N content was lower in infected hosts, and all other comparisons between uninfected and infect host elemental content were not significant (p > 0.05). Asterisk denotes significant differences in host stoichiometry. Filled circles are group means and unfilled circles are the raw data that are jittered to show their distribution, and error bars are standard errors.

**Figure S2.**
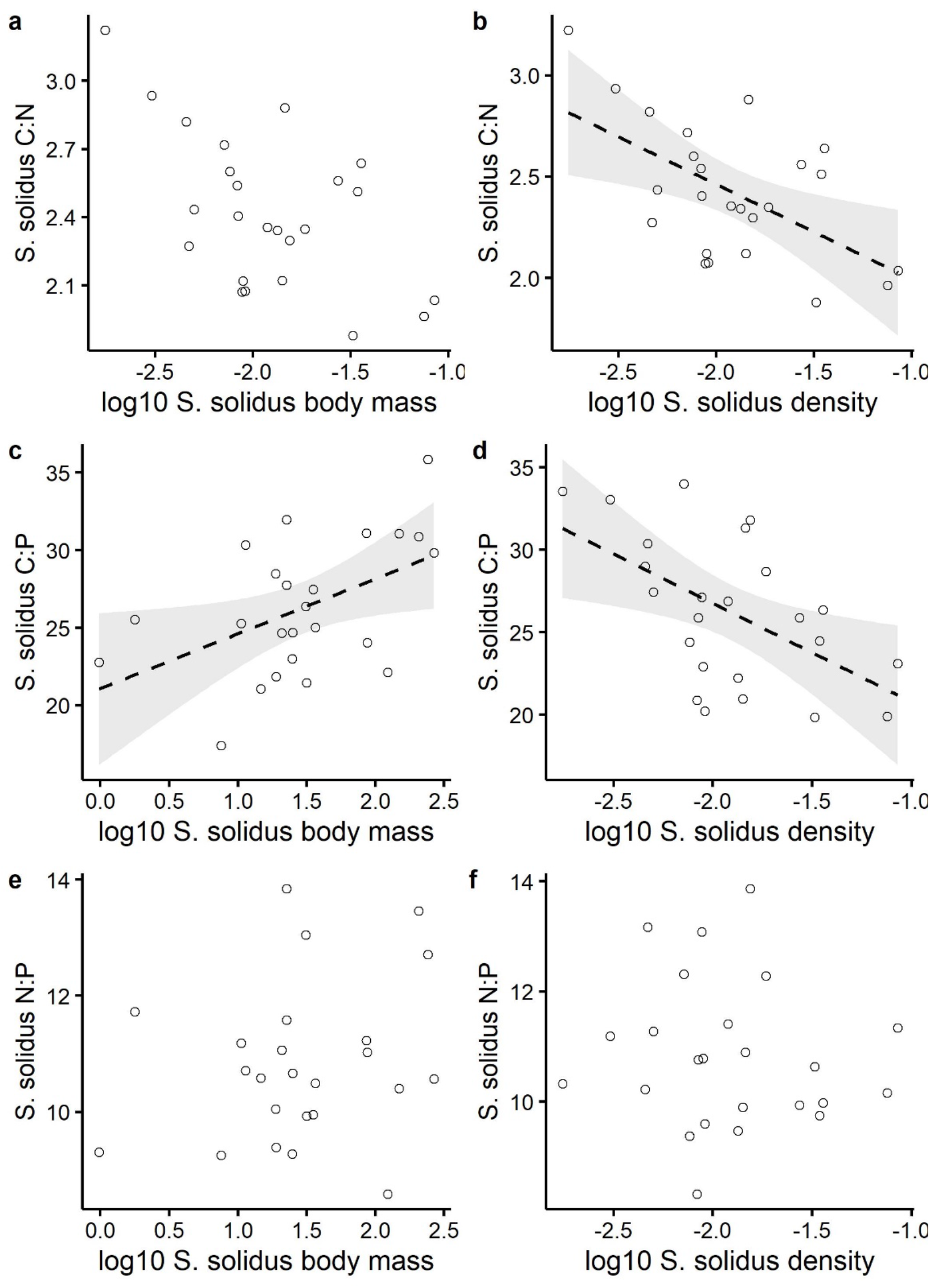
Parasite C:N, C:P, and N:P showed more variable relationships with parasite body mass and parasite density. Plotted are the partial residuals of the statistical model for parasite C:N (a-b), C:P (c-d), and N:P (e-f). Dashed lines are the estimated model fits and grey shaded area is the 95% confidence interval.

